# A mathematical investigation of the interplay between vasculature and intratumoral cellular heterogeneity during tumor progression

**DOI:** 10.64898/2026.08.26.747242

**Authors:** Supriyo Ghosh, Gopinath Sadhu, D C Dalal

## Abstract

Tumors consist of heterogeneous phenotypic cells, such as normoxic cells, which are highly proliferative, and hypoxic cells, which are less proliferative. Their phenotypic switching depends on tumor microenvironmental factors, such as oxygen and nutrient concentrations supplied by local blood vessels. However, during ongoing angiogenesis, the process of sprouting new blood vessels at the tumor site from pre-existing blood vessels, and how this phenotypic switching affects and impacts tumor growth, remains poorly understood. In this article, we formulate a mathematical model to elucidate the crosstalk between vasculature and tumor cellular heterogeneity during tumor progression. The model results show a strong agreement with the experimental data. Our simulation results demonstrate that ongoing angiogenesis increases tumor growth rate. In addition, we observe that the influence of hypoxic cells on phenotypic switching from normoxic to hypoxic is more pronounced than their influence on the transition from hypoxic to normoxic. Furthermore, we perform a global sensitivity analysis using the Sobol’s method to assess the importance of the model’s parameters. It highlights that the volume at which blood vessels attain half-maximal rate has the maximum effect on the model.

## 1 Introduction

Tumor growth is a complex biological phenomenon. The development of a tumor begins with a genetically altered cell that acquires the ability to proliferate indefinitely, uncontrollably, and rapidly [1]. As a result, a cluster of cells is formed, known as an avascular tumor, whose nutrients and oxygen are supplied by the existing primary blood vessels. In the initial stage, the cells are well nourished, well oxygenated, and highly proliferative. These cells are referred to as normoxic cells. Due to the high oxygen consumption at the tumor periphery and the increasing diffusion distance between the primary blood vessels and the tumor site, an oxygen gradient gradually develops [2]. Consequently, the normoxic cells in the tumor core experience oxygen stress and shift their metabolism from oxidative phosphorylation to glycolysis [3]. Simultaneously, they undergo a phenotypic transition from the normoxic to the hypoxic state, during which they become less proliferative or cease proliferating. Furthermore, when hypoxic cells are subjected to prolonged oxygen deprivation, they eventually die, leading to the formation of a necrotic region at the tumor core [4]. Experimental studies indicate that avascular tumors cannot grow beyond a size of approximately (1–2 *mm*^3^) [5]. To support further growth, tumors develop their own vascular network within their territory through a process known as angiogenesis [6]. Hypoxic cells play an important role in angiogenesis by secreting vascular endothelial growth factor (VEGF), which acts as a chemotactic signal, stimulating the growth of blood vessels toward the tumor site [7]. Once new blood vessels are established, the tumor microenvironment becomes enriched with nutrients and oxygen, enabling hypoxic cells to revert to the normoxic phenotype [8, 9]. However, our understanding of how ongoing vasculature influences phenotypic heterogeneity within tumors remains limited.

Mathematical models serve as excellent substitutes for time-consuming and expensive experimental studies of biological phenomena, especially cancer [10–12]. In the literature, various modeling frameworks are broadly categorized into three types: continuum, discrete, and hybrid [4]. In the continuum modeling framework, biological quantities are treated as continuous variables, and the underlying phenomena are described using ordinary differential equations (ODEs) and partial differential equations (PDEs). On the other hand, discrete modeling approaches treat each cell in the tumor tissue as an individual entity, enabling the capture of cell-cell interaction dynamics through agent-based models (ABM) and cellular Potts models (CPM)[13]. Hybrid modeling frameworks effectively capture the multiscale nature of biological phenomena by combining continuum and discrete modeling approaches. Numerous continuum Mathematical modeling approaches have been proposed to capture both avascular [14, 15] and vascular tumor growth dynamics [16, 17]. In avascular tumor models, blood vessel development is not considered; instead, blood vessel density is assumed to be constant [18, 19]. On the other hand, for vascular tumor models, the dynamic behavior of endothelial cells of blood vessels, such as proliferation and death, is accounted for. In the avascular phase, the tumor consumes oxygen and nutrients, which are supplied from nearby existing blood vessels, and begins to proliferate [20, 21]. As the tumor grows, the demand for nutrients and oxygen increases rapidly, and the pre-existing blood vessels are unable to meet this demand; therefore, an oxygen gradient forms. Consequently, oxygen heterogeneity occurs within the tumor domain. To meet the demand for oxygen, tumors induce angiogenesis [22] and establish a vascular network within the tumor ecosystem. Stamper et al. [23] provided a mathematical model of vascular tumor growth, in which both angiogenesis and vasculogenesis contribute to the formation of blood vessels. Due to vascular instability, the vascular tumor exhibits a heterogeneous cellular phenotype. Villa et al. [24] developed a phenotype-structured model to investigate phenotypic heterogeneity in vascularized tumors; however, their model did not explicitly account for ongoing angiogenesis. Recently, Borzouei et al. [25] developed a two-dimensional hybrid modeling framework to investigate both phenotypic heterogeneity and angiogenesis. They found that angiogenesis does not uniformly benefit the whole tumor. Instead, cells located near blood vessels proliferate, while a substantial population of dead cells accumulates in the tumor core. But their model did not address the interactions among heterogeneous tumor cells. The present study aims to investigate tumor cell heterogeneity during angiogenesis and assess how heterogeneous tumor cells influence one another.

In this article, we present a mathematical model to illustrate cellular heterogeneity in tumors during angiogenesis and the cross-talk among heterogeneous tumor cells in the presence of blood vessels. Our model illustrates the temporal evolution of the volumes of normoxic, hypoxic, and necrotic cells, as well as blood vessels. We assume that phenotypic switching among cells depends on the local blood vessel density. Previous studies reported that mesenchymal cells, which are hypoxic, resist phenotypic switching from the mesenchymal to the epithelial (normoxic) state, and that hypoxic cells promote the transition from normoxic to hypoxic cells [26]. We incorporate these biological phenomena into our model to explore how hypoxic cells influence cellular heterogeneity during ongoing angiogenesis. The model simulation shows excellent agreement with experimental data from a mouse model. In addition, our simulation results demonstrated that ongoing angiogenesis affects normoxic cells to such an extent that, despite a decrease in the volume of hypoxic and necrotic cells, total tumor volume remains elevated. Moreover, we employ Sobol’s variance method to assess the model’s global sensitivity in the parameter space. Sensitivity analysis indicates that the volume at which blood vessels attain it’s half-maximal proliferation rate is the most sensitive, whereas the degradation rates of normoxic cells, necrotic cells, and angiogenic threshold, as well as the influence of hypoxic cells on phenotypic switching from normoxic to hypoxic and from hypoxic to normoxic, are less sensitive.

## 2 Mathematical model

Here, we propose a mechanistic model to study intratumoral heterogeneity during ongoing angiogenesis, accounting for the influence of hypoxic cells on phenotypic switching between normoxic and hypoxic states. Our model consists of four components: normoxic cells volume (*n*(*t*)), hypoxic cells volume (*h*(*t*)), necrotic cells volume (*s*(*t*)), and blood vessels volume (*v*(*t*)), where *t* is time. The unit of the volume is considered as *mm*^3^. Our model is based on the following biological underpinnings:

- Hypoxic cells elevate the transition from normoxic to hypoxic.
- Hypoxic cells resist the switch from hypoxic to normoxic.
- Proliferation of normoxic cells increases in the presence of blood vessels.
- Phenotypic switching from normoxic to hypoxic state is reduced in the presence of blood vessels.
- Phenotypic switching from hypoxic to normoxic state increases in the presence of blood vessels.

### 2.1 Normoxic cell (*n*) dynamics

We consider that normoxic cells proliferate at a rate *λ* in a logistic manner, and that their proliferation depends on local blood vessel density. Normoxic cells change their phenotype to hypoxic cells [27] at a rate *µ*_1_, and the reverse transition, i.e., switching from hypoxic to normoxic [27], occurs at a rate *µ*_2_. Normoxic cells switch to necrotic cells at a rate *δ*_1_. The temporal dynamics of normoxic cells are given by,

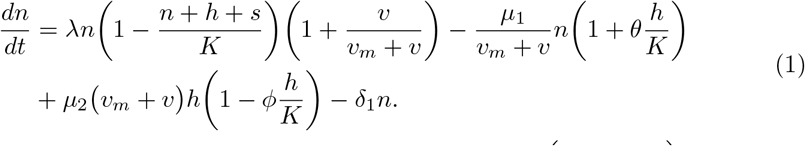

Here, the first term of the right-hand side (RHS) of Eq. (1), 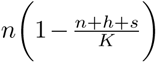, represents proliferation of normoxic cells [28], where *K* is the carrying capacity of the tumor. As the proliferation of normoxic cells increases in the presence of blood vessels, so we incorporated the term 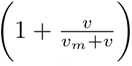, which signifies the effects of blood vessels in the proliferation of normoxic cells, where *v_m_* denotes the half-saturation rate. Even for *v* = 0, normoxic cells proliferates logistically. Blood vessel volume amplifies the proliferation of normoxic cells by the term 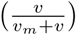, which is assumed to be a form of Michaelis-Menten kinetics. The second term of the RHS of Eq. (1) represents normoxic cells switching to hypoxic cells at a rate *µ*_1_ when the local blood vessels volume decreases which is represented by the term 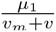, and this switching rate is amplified in the presence of hypoxic cells. Hypoxic cells promote the transition from normoxic to hypoxic i.e. it promote normoxic cells to become hypoxic, this phenomena is mathematically represented by the term 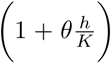, where *θ* is the promoting influence parameter, which signifies the influence of hypoxic cells in phenotypic switching from normoxic to hypoxic. The third term on the RHS of Eq. (1) denotes that hypoxic cells adopt a normoxic phenotype at a rate *µ*_2_, which increases with local blood vessel density, and this is model as *µ*_2_(*v_m_* + *v*). This phenotype adaption from hypoxic to normoxic slows down in the presence of local hypoxic cells, i.e. hypoxic cells resists itself to become normoxic again and this phenomena is represented mathematically by the term 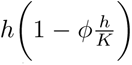, where *ϕ* is the resisting influence parameter that signifies the influence of hypoxic cells in phenotypic switching from hypoxic to normoxic state. The last term indicates that the normoxic cells die at a rate *δ*_1_. All the parameters are summarized in the table 1.

**Table 1.** Parameter description and values.

| Parameter | Description | Dimension | Dimensional value | Non-dimensional value | Reference |
| --- | --- | --- | --- | --- | --- |
| $\lambda$ | Proliferation rate of normoxic cells | $day^{-1}$ | 0.215 | $1.1 \times 10^{-2}$ | estimated |
| $K$ | Carring capacity | $mm^3$ | 195 | - | [34] |
| $\delta_1$ | Conversion of normoxic to necrotic | $day^{-1}$ | $4.0 \times 10^{-10}$ | $1.8 \times 10^{-6}$ | [35] |
| $\delta_2$ | Conversion of hypoxic to necrotic | $day^{-1}$ | $3.8 \times 10^{-6}$ | $1.7 \times 10^{-2}$ | [35] |
| $\nu$ | Degradation of dead cell | $day^{-1}$ | $\frac{\lambda}{10}$ | $1.1 \times 10^{-3}$ | assumed |
| $\gamma$ | Proliferation of endothelial cell | $day^{-1}$ | $\frac{\lambda}{3}$ | $3.7 \times 10^{-3}$ | assumed |
| $\beta$ | Degradation of blood vessel | $day^{-1}$ | 0 | 0 | assumed |
| $\mu_1$ | Conversion of normoxic to hypoxic | $mm^3 day^{-1}$ | 3.8 | $10^{-2}$ | estimated |
| $\mu_2$ | Conversion of hypoxic to normoxic | $mm^{-3} day^{-1}$ | 0.1 | - | [33] |
| $v_m$ | Half cycle | $mm^3$ | 0.1 | $1.02 \times 10^{-1}$ | - |
| $\theta$ | Influence of hypoxic cells in phenotypic switching from normoxic to hypoxic | - | - | 1 | - |
| $\phi$ | Influence of hypoxic cells in phenotypic switching from hypoxic to normoxic | - | - | 1 | - |

### 2.2 Hypoxic cell (*h*) dynamics

Hypoxic cells are reported as non-proliferating [29–31]. Therefore, any growth term in hypoxic cell dynamics is not considered. Hypoxic cells are formed only from the transition from normoxic cells. The temporal evolution of hypoxic cell dynamics is modeled as,

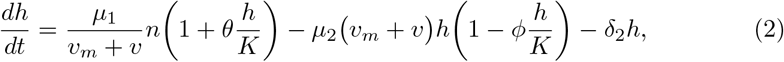

the first term of the RHS of Eq. (2) represents normoxic cells switching to hypoxic cells when local blood vessel density decreases, and the second term denotes that the hypoxic cells revert to normoxic cells when local blood vessels increase. Last term highlights the death of hypoxic cells at a rate *δ*_2_.

### 2.3 Necrotic cell (*s*) dynamics

The temporal dynamics of necrotic cells are given by,

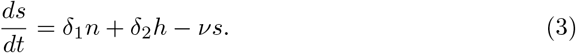

Here, *δ*_1_*n* is the conversion of normoxic cells to necrotic cells, *δ*_2_*h* is the conversion of hypoxic cells to necrotic cells and *ν* denotes the rate of natural decay of necrotic core.

### 2.4 Blood vessel (*v*) dynamics

The growth of blood vessels is necessary for a tumor to grow. After a certain volume, say *T*_0_, the inner core of the tumor becomes hypoxic, and hypoxia triggers the growth of blood vessels towards the tumor. We assume that blood vessels proliferate logistically [17] and undergo a natural decay at a rate *ν*. All these process put in mathematical form as,

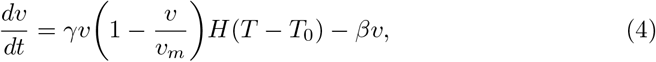

where *γ* is the proliferation rate of blood vessels and *H*(*x*) is the Heaviside function and define as

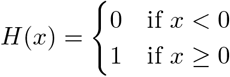

and *T* = *n* + *h* + *s*, the total tumor volume.

### 2.5 Initial conditions

In order to solve our model uniquely, we need to specify the initial conditions. Initially, we consider that the tumor consists only of normoxic cells. When normoxic cells begin to proliferate, they consume nutrients and oxygen from nearby blood vessels, thereby creating a hypoxic tumor microenvironment. Hypoxic cells then arise, so initially, no hypoxic or necrotic cells are present in the tumor. We assume that at the initial time (*t* = 0) the normoxic cell volume is *n*_0_ and the blood vessel volume is *v*_0_. Hence, the initial conditions are

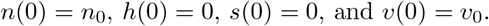

## 3 Non-dimensionalization

Let the carrying capacity *K* be the characteristic volume and 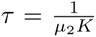 be the characteristic time, which is the timescale of hypoxic to normoxic phenotype. We use the following transformation to non-dimensionalize the governing equations.

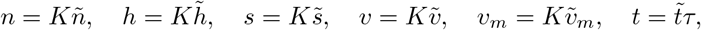

where tilde sign denotes the non-dimensionalize unit. The corresponding non-dimensional parameters are: 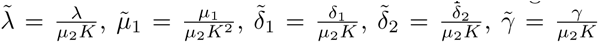, 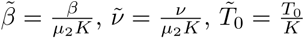

For compactness of notation, we drop the tilde sign from variables and parameters in the non-dimensional model. Thus, the dimensionless governing equations become

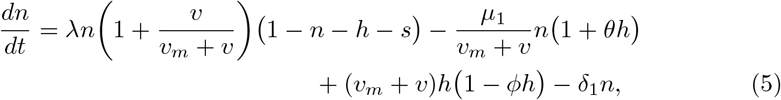

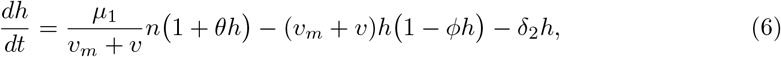

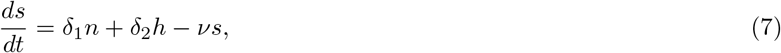

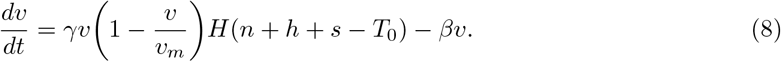

The non-dimensionalized initial conditions are

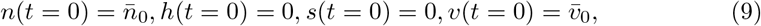

where 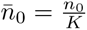 and 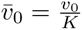.

## 4 Parameter choice

In order to reduce the complexity of the model, we assume that 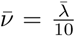 and 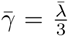. Default values of both *θ* and *ϕ* are taken as 1. We estimate the parameter 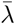 and 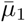 from the data set given in [32]. We use the value of *µ*_2_ = 0.45 *day^−^*^1^ which lies in the range as suggested by Wenxiang Liu [33]. In order to match the unit of *µ*_2_, we have divided *µ*_2_ by a reference volume. Here, we consider the reference volume to be 4.4 *mm*^3^, which we have taken as the geometric mean of the carrying capacity and the initial normoxic cell volume. All the parameters are used in the model, and their corresponding non-dimensional values are shown in table 1.

## 5 Results

We solve our non-dimensionalized model, Eqs. (5) -(8) with the initial conditions Eq. (9) and *n̄*_0_ = 0.123, *h̄*_0_ = 0, *s̄*_0_ = 0, *v̄*_0_ = 0.0109. The values of the parameters are taken from the table 1. We have used the ODE45 solver in MATLAB software to solve the governing equations.

### 5.1 Model validation

In this section, the proposed model is validated against experimental data provided by Malekian et al. [32]. They showed that the expression of angiogenic factors occurs in various stages of a Triple-Negative Breast Cancer(TNBC). After injecting 4T1 breast cancer cells into mice to generate TNBC tumors, they separate the tumors by volume and calculate the proportions of angiogenic markers. They found out that angiogenic-related markers begin to increase at the very early stages of the tumor, even when the tumor volume is less than 100*mm*^3^. To compare our model with the experimental data, we used the parameter values from table 1, except for the carrying capacity, which is set to *K* = 2000. Since the experimental paper reports a total tumor volume of approximately 2000 *mm*^3^, *K* = 2000 is justified. We determine the total tumor volume (*n* + *h* + *s*) in our model and compare it with the experimental data [32]. Our simulation results agree perfectly with the experimental data (figure 2). This ensures the reliability of our model.

**Fig. 1.**
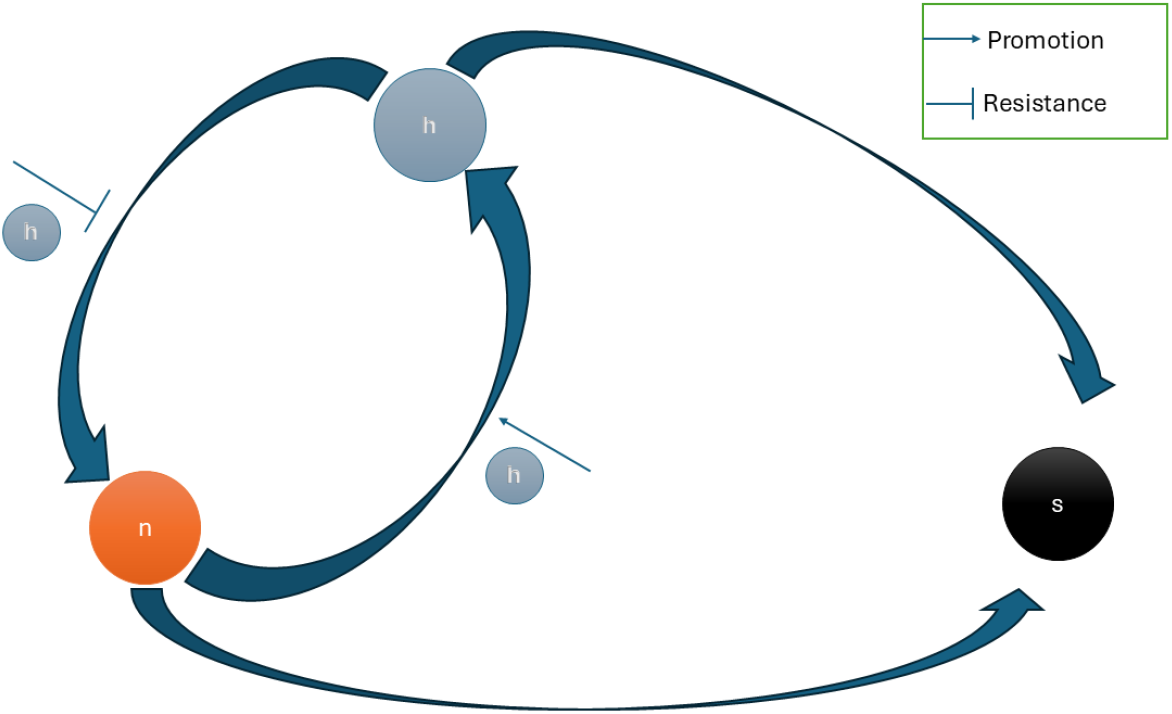
Schematic of normoxic(n), hypoxic(h), and necrotic(s) cells showing that hypoxic cells promote the transition from normoxic to hypoxic, and they resist the transition from hypoxic to normoxic.

**Fig. 2.**
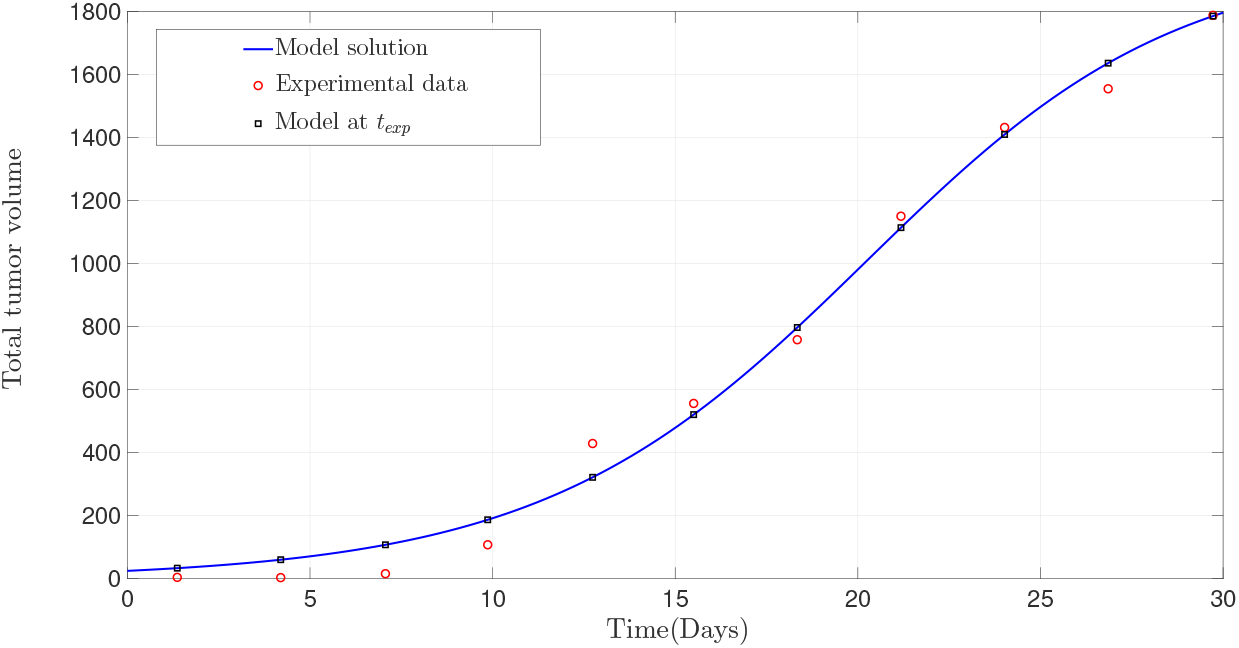
Comparison of simulated results with the data provided by Malekian et al.[32]

### 5.2 Effects of vasculature on tumor growth dynamics

In this section, we explore how tumor growth dynamics change over time during ongoing vasculature. Figure 3 displays the temporal evolution of normoxic, hypoxic, necrotic cells and blood vessels, and the total volume of the tumor. We observe that the volume of normoxic cells increases rapidly during the initial transient period and then reaches a steady state. On the other hand, it has been observed that hypoxic cells increase in number during the initial stage of growth. However, hypoxic cells also attain a steady state subsequently. It is interesting to note that hypoxic cells take longer than normoxic cells to reach a steady state. It occurs because the tumor initially consists predominantly of normoxic cells, which are highly proliferative, and they switch to a hypoxic phenotype. As the tumor reaches a critical mass *T*_0_, the angiogenic progression is triggered. We observe that once the blood vessels stabilize in the system, the hypoxic core reaches a steady state. Therefore, it can be concluded that the lag time required for hypoxic cells to attain a steady state in the tumor system depends on the stabilization of the local blood vessels.

**Fig. 3.**
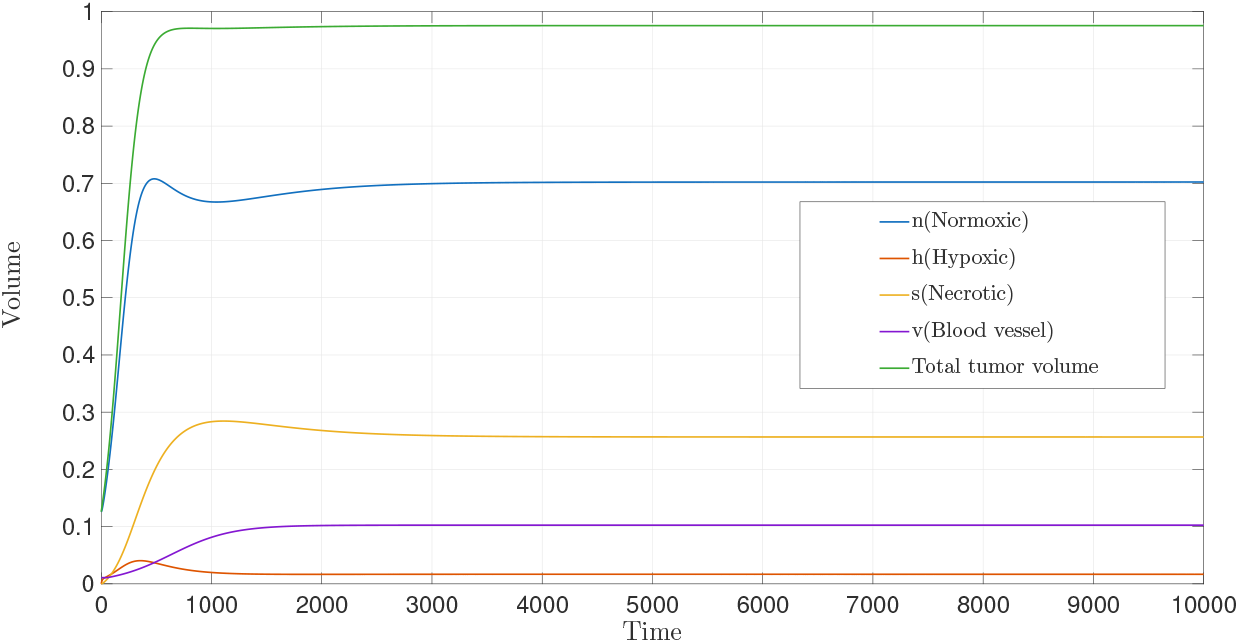
Temporal evolution of normoxic, hypoxic, necrotic cell volume, blood vessel volume and total tumor volume.

### 5.3 Crosstalk between ongoing angiogenesis and intracellular heterogeneity

In this section, we investigate the crosstalk between vasculature and intracellular heterogeneity during tumor growth. Normoxic cells are closest to blood vessels, so they receive adequate oxygen after angiogenesis, which leads to high proliferation ability. Consequently, normoxic cell volume increases in vascular tumors compared to that in avascular tumors (Figure 4a). Once adequate blood vessels are available in the tumor microenvironment, the transition from normoxic to hypoxic decreases, while the reverse transition from hypoxic to normoxic increases. As a result, the volume of hypoxic cells decreases after angiogenesis (Figure 4b). Necrotic cell volume also follows a similar pattern as of hypoxic cell volume, as the transition rate *δ*_2_ is higher than *δ*_1_. In ongoing angiogenesis, the predominance of normoxic cell volume leads to an increase in total tumor volume despite a decline in hypoxic and necrotic cell volume (Figure 4).

**Fig. 4.**
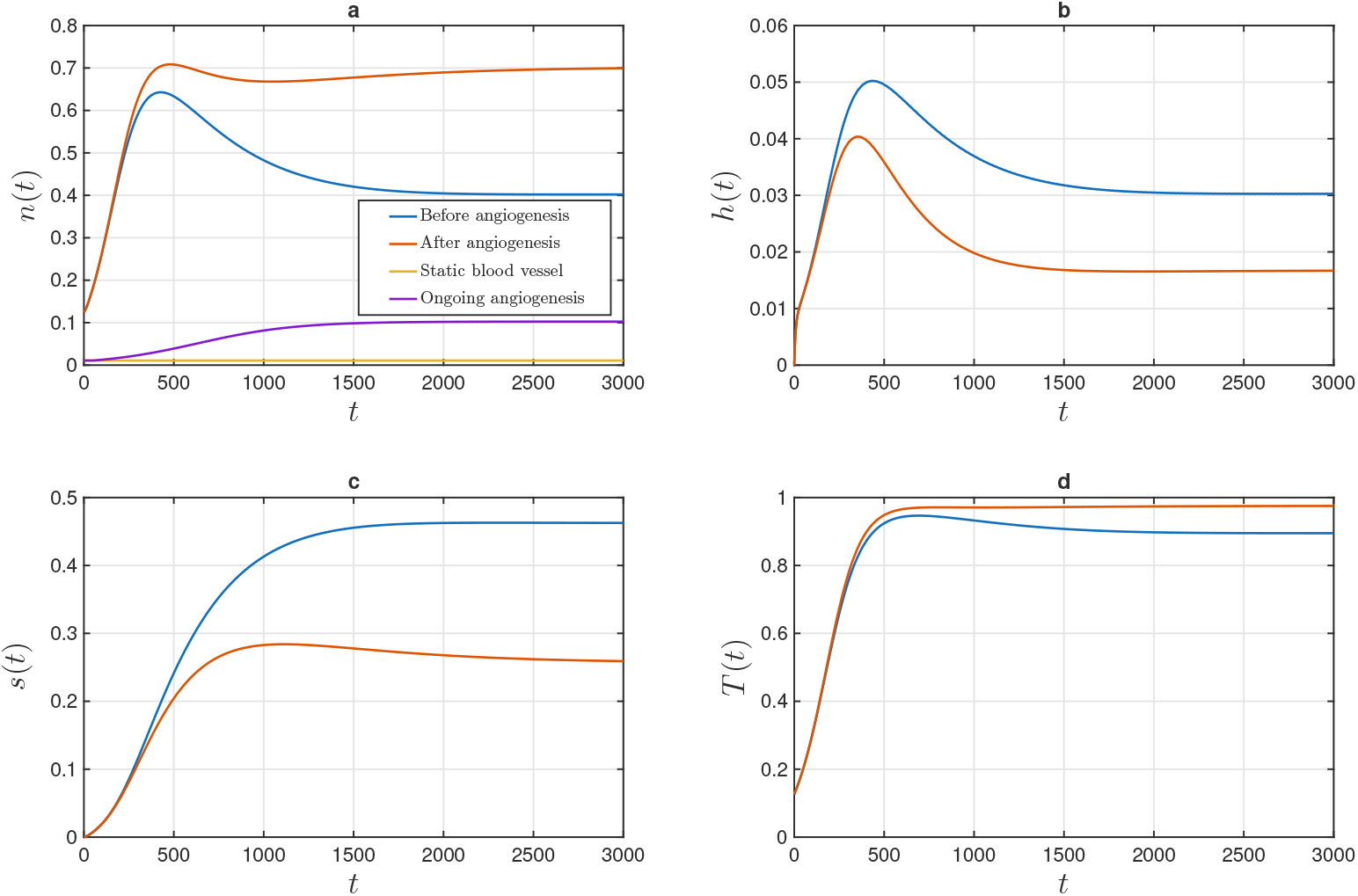
Effects of angiogenesis on (a) normoxic cell volume, (b) hypoxic cell volume, (c) necrotic cell volume, (d) total tumor volume.

From this section, we conclude that the effects of angiogenesis primarily affect normoxic cells, whereas hypoxic and necrotic cells respond negatively to post-angiogenesis. Because normoxic cells are closest to the blood vessels, they benefit the most.

### 5.4 Effects of the promoting coefficient *θ* and the resisting coefficient *ϕ* on normoxic, hypoxic, necrotic cells and blood vessels

It is observed that hypoxic cells promote normoxic cells to switch into hypoxic phenotype [36]. Here, we aim to visualize cross-talk between normoxic and hypoxic cells, that is, the impact of hypoxic cells on normoxic cells, in the presence of blood vessels. So, we run a series of simulations to examine the influence of *θ*.

Figure 5a demonstrates that during ongoing angiogenesis, normoxic cell volume decreases as *θ* goes up from 0 to 7. This decline occurs because hypoxic cells promote the conversion of normoxic cells into hypoxic cells. As the promoting coefficient *θ* increases, a larger proportion of normoxic cells switch to hypoxic cells, resulting in a reduction in normoxic cell volume. In contrast, hypoxic cell volume (figure 5b) increases with rising *θ*, as the promotion of normoxic to hypoxic cells becomes more significant. Necrotic cell volume follows a similar pattern (figure 5c) as hypoxic cell volume, since the transition rate from hypoxic to necrotic cells is higher compared to the rate from normoxic to necrotic cells. Static blood vessels display a comparable pattern to ongoing angiogenesis for the same reason stated above. In the blood vessel profile (figure 5d), when the total tumor volume exceeds the threshold *T*_0_, blood vessels begin to proliferate. It is observed that, for *θ* = 7, the total tumor volume surpasses the threshold *T*_0_ earlier than it does for other values of *θ*. Also, the influence of *θ* is less significant in the blood vessels profile compared to other cell compositions.

**Fig. 5.**
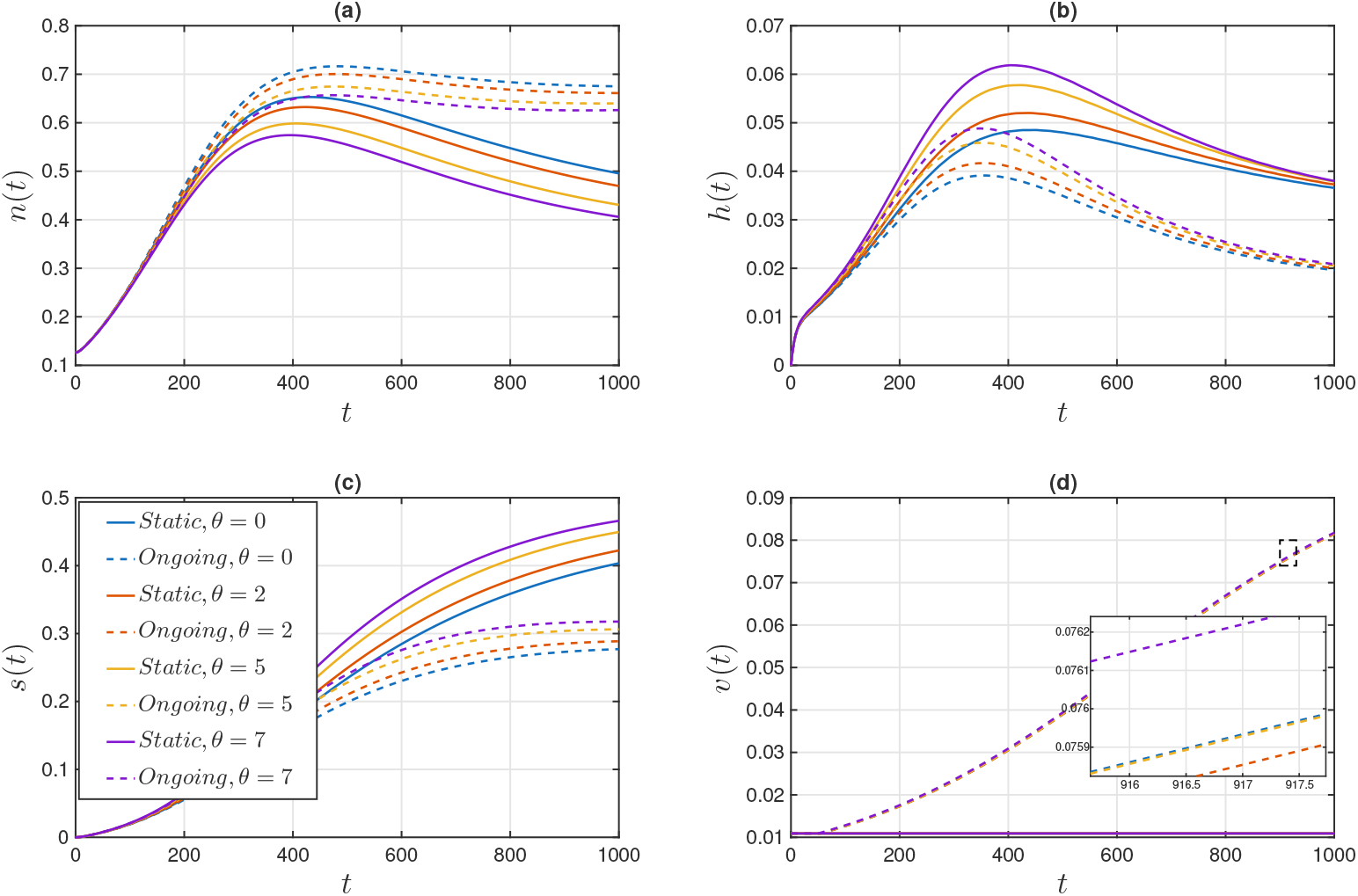
Time evolution simulation of (a) normoxic cell volume, (b) hypoxic cell volume, (c) necrotic cell volume, (d) blood vessel volume, for different values of *φ*, fixing *θ* = 1 and *T*_0_ = 0.2. The dotted line is the volume profile, including angiogenesis, i.e., when the blood vessel is non-constant, and the solid line is the volume profile excluding angiogenesis, i.e., when the blood vessel is static.

Hypoxic cells retain a hypoxic memory even after receiving adequate oxygen and nutrients, suggesting that hypoxic cells resist switching to a normoxic phenotype [37]. Here, we conduct a series of simulations varying *ϕ*to observe its effects on each cell type and blood vessels. In the case of ongoing angiogenesis, the fraction of normoxic cells decreases as *ϕ* increases from 0 to 1 (figure 6a). This is because hypoxic cells resist switching to normoxic cells; as the resistance coefficient increases, normoxic cell volume decreases, while hypoxic cell volume increases (figure 6b). Necrotic cell volume also follows a similar pattern as hypoxic cell volume, since the transition from hypoxic to necrotic is higher than from normoxic to necrotic (figure 6c). Static blood vessels exhibit a pattern similar to that observed in ongoing angiogenesis for the same reason discussed above. For the blood vessel profile, we see that for *ϕ* = 0, the total tumor volume crosses the threshold *T*_0_ earlier than that for other values of *ϕ* (figure 6d).

**Fig. 6.**
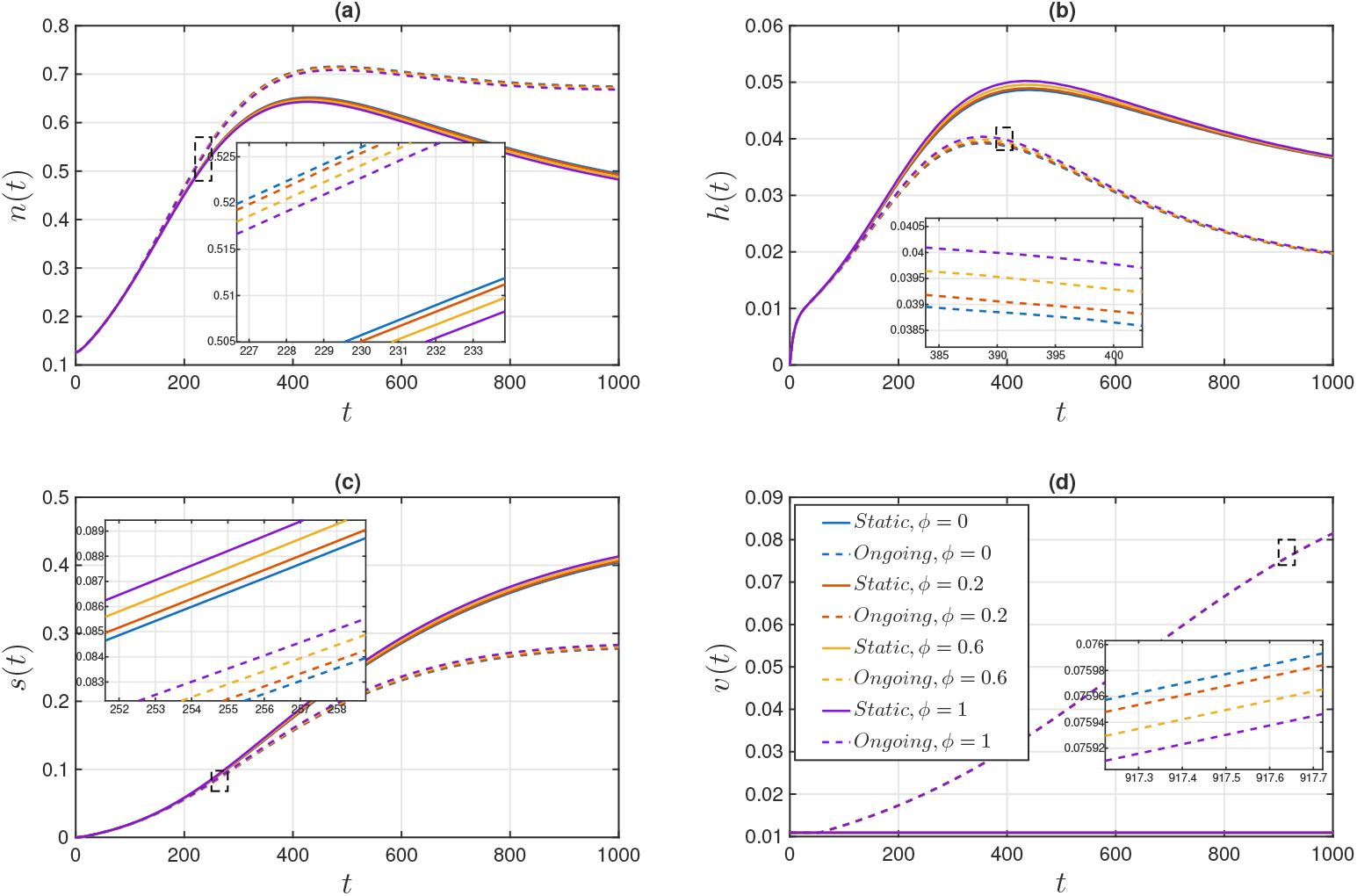
Time evolution simulation of (a) normoxic cell volume, (b) hypoxic cell volume, (c) necrotic cell volume, (d) blood vessel volume, for different values of *ϕ*, fixing *θ* = 1 and *T*_0_ = 0.2. The dotted line is the volume profile, including angiogenesis, i.e., when the blood vessel is non-constant, and the solid line is the volume profile excluding angiogenesis, i.e., when the blood vessel is static.

Clearly, from figures 5 and 6, we can notice that the effects of *θ* are higher than that of *ϕ*, i.e., the effects of hypoxic cells in the transition from normoxic to hypoxic are higher than the transition from hypoxic to normoxic.

### 5.5 Volume profile of normoxic, hypoxic and necrotic cells for the variation of angiogenic threshold *T*_0_

Tumor volume is highly influenced by the angiogenic threshold *T*_0_. When tumor volume surpasses a specific size, the tumor core becomes hypoxic. The hypoxic core subsequently releases pro-angiogenic factors such as VEGF and FGF, which are critical in initiating the angiogenic switch [38]. After the switch, blood vessels begin to proliferate. As shown in figure 7a, during ongoing angiogenesis, the volume of normoxic cells decreases as the threshold *T*_0_ increases from 0.1 to 0.9. A lower *T*_0_ value indicates that blood vessel proliferation commences earlier, thereby increasing the availability of nutrients and oxygen to normoxic cells. Consequently, the volume of normoxic cells is higher at a lower angiogenic threshold. Early access to oxygen and nutrients reduces the transition from normoxic to hypoxic conditions, leading to a decrease in hypoxic cell volume at lower angiogenic thresholds (figure 7b). As the threshold increases, angiogenesis is delayed (figure 7c), resulting in poor vascularization of the tumor and an increase in necrotic cell volume. Blood vessel volume increases as *T*_0_ decreases (figure 7d). Importantly, static blood vessels do not influence normoxic, hypoxic, or necrotic cells. Therefore, all values of *T*_0_ coincide for the static case.

**Fig. 7.**
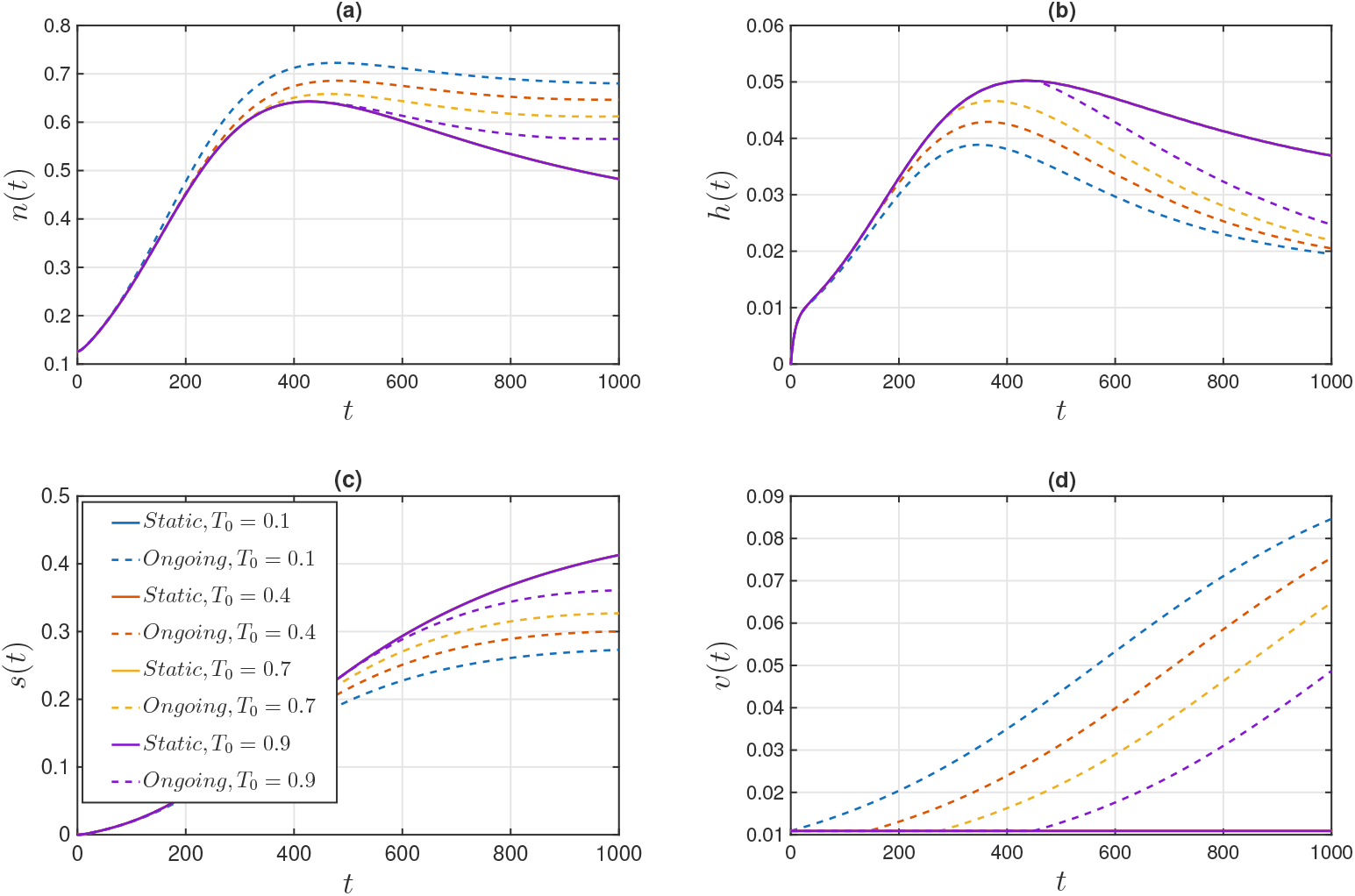
Time evolution simulation of (a) normoxic cell volume, (b) hypoxic cell volume, (c) necrotic cell volume, (d) blood vessel volume, for different values of *T*_0_, fixing *θ* = 1 and *ϕ* = 1. The dotted line is the volume profile, including angiogenesis, i.e., when the blood vessel is non-constant, and the solid line is the volume profile excluding angiogenesis, i.e., when the blood vessel is static.

## 6 Sensitivity analysis

Sensitivity analysis is performed to understand how changes in the model parameters would influence the output. It can primarily be done in two ways: (1) local sensitivity analysis, (2) global sensitivity analysis. Local sensitivity analysis examines the effect of a single parameter, keeping all others fixed, whereas global sensitivity analysis allows parameters to vary simultaneously across the parameter space. Our aim is to determine which parameter would significantly influence total tumor volume. To understand the sensitivity of the parameters, global sensitivity analysis is performed. We use the Sobol method for global sensitivity analysis [39, 40].

Let *Y* = *f* (*X*_1_*, X*_2_*, . . ., X_k_*), where *X*_1_*, X*_2_*, . . ., X_k_* are input parameters, and *Y* is the output. The variance of *Y* is given by *V* (*Y* ).

The first-order sensitivity index is given by

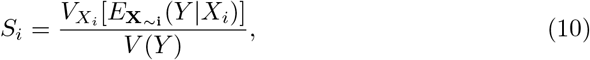

where *E***_X_***_∼_***_i_** (*Y |X_i_*) is the mean of the argument *Y* taken over all the factors but *X_i_*. *V_Xi_* [*E***_X_***_∼_***_i_** (*Y |X_i_*)] is the variance of *E***_X_***_∼_***_i_** (*Y |X_i_*) taken over *X_i_*. The first-order sensitivity index *S_i_* represents the direct effects of each parameter on the variance. While the total effect index measures the total effect of *X_i_* on the variation caused by the first and higher order effects due to interactions, it is given by

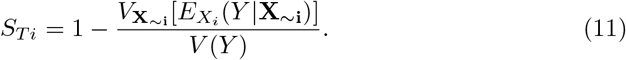

In this sensitivity analysis, we consider the parameters from Table 1 (non-dimensional values), assuming a uniform distribution spanning 0.9 to 1.1 times their specified values. The output is the total tumor volume at *t* = 2000, where the steady state is attained. Within the specified variations, the order of the parameters that contribute to the output is *v_m_ > µ > λ > δ*_2_ *> γ > δ*_1_ = *ν* = *θ* = *ϕ* = *T*_0_. We observe that the significances of *θ*, *ϕ*, and *T*_0_ are very low. This low impact is because only a narrow range of variation (0.9 to 1.1 times the specified value) was used for these parameters, which limits the extent of their influence on the total tumor volume output.

## 7 Discussion

Intratumoral heterogeneity is a major barrier to anticancer therapy [41], and blood vessels are central contributors to this heterogeneity. Sprouting vessels formed during angiogenesis are leaky and fragile in nature. As a result, the distributions of nutrients and oxygen become spatially heterogeneous, leading to the emergence of intracellular phenotypic heterogeneity within the tumor, such as normoxic and hypoxic phenotypes [9, 26]. These hypoxic and normoxic cells switch their phenotypes depending on the local nutrient concentration. Experimental studies have also reported that hypoxic cells promote phenotypic switching from the normoxic to the hypoxic state while resisting the reverse transition to the normoxic state. However, our understanding of how ongoing vasculature influences intratumoral phenotypic heterogeneity and, consequently, tumor growth remains limited. In this study, we have formulated a mathematical model to examine the phenotypic heterogeneity of tumor cells during ongoing angiogenesis and examined how interactions among heterogeneous cell populations influence tumor growth dynamics. We have found that our model simulation results are in excellent agreement with the experimental data of Malekian et al. [32] (Figure 2).

Our findings suggest that ongoing angiogenesis mostly affects normoxic cells. In addition, we have observed that after angiogenesis is initiated, normoxic cells increase rapidly, while hypoxic cells decline below (figure 4). A similar observation was made in the experimental study by Ghajar et al. [42]. They found that, in breast cancer cells, stable vasculature supports a dormant state, whereas sprouting vessels accelerate tumor growth. Additionally, our model reveals that the influence of hypoxic cells in promoting normoxic cells into hypoxic cells is greater than the influence of hypoxic cells in resisting itself to become normoxic cells [figure 5, 6]. From figure 7, we observe that the angiogenic threshold is meaningless in the static case, whereas ongoing angiogenesis affects all the heterogeneous tumor cells significantly. Normoxic cells respond negatively to increasing threshold, while hypoxic and necrotic cells respond positively. We provide an analytical solution with two different biological scenarios. Firstly, when the tumor is fully vascularized (assuming *v → v_∞_*, Appendix A.1) and secondly, when the tumor is poorly vascularized (assuming *v →* 0, Appendix A.2). For the first case, we compare the analytical solution with our numerical solution, and we see that the analytical solution exactly matches the numerical solution (figure 9).

**Fig. 8.**
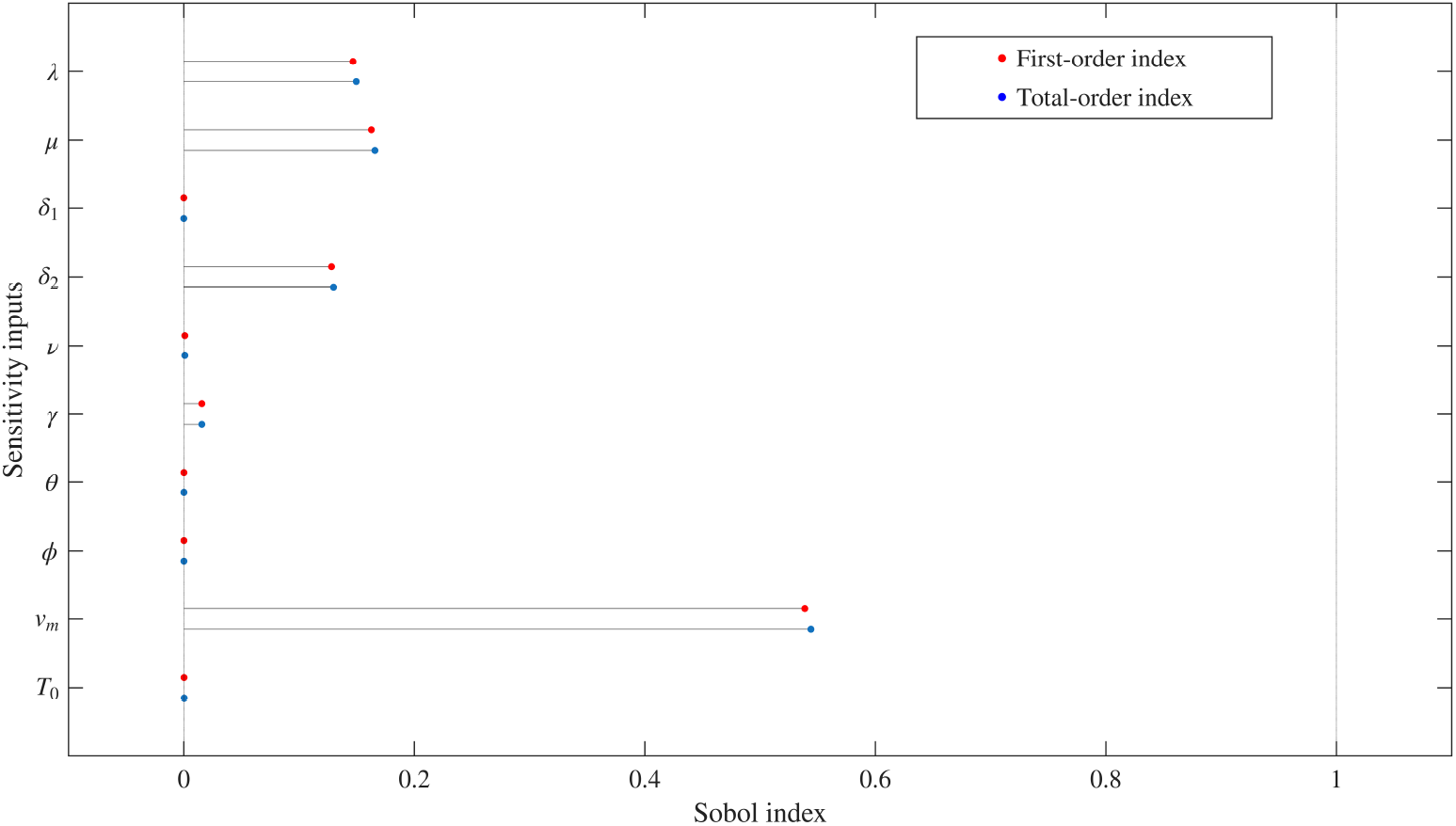
Sensitivity analysis of all parameters, with the output being total tumor volume, was evaluated at time *t* = 2000, when the steady state is attained.

**Fig. 9.**
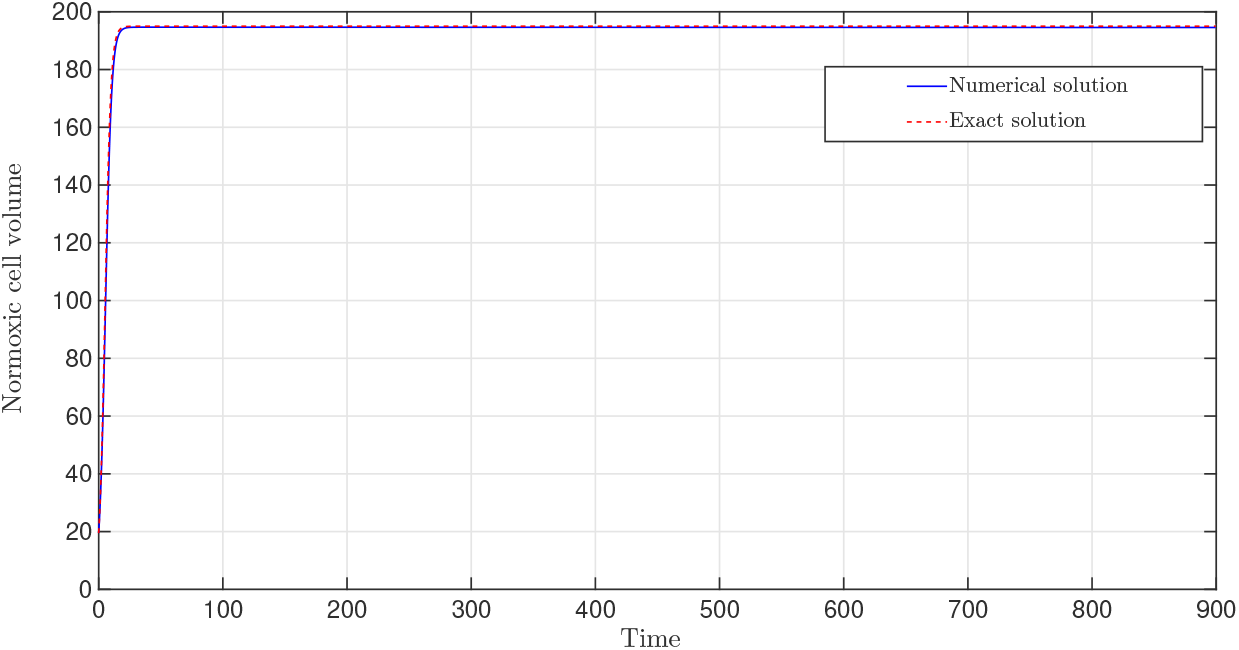
Comparison between numerical simulation and exact solution, which is obtained in Eq. (13).

Our model investigates only the temporal dynamics of tumor growth and the corresponding intratumoral cellular heterogeneity in avascular and ongoing angiogenic tumors. However, these dynamics also depend on spatial properties, such as the spatial distribution of blood vessels, where tumor cell proliferation rates increase in regions of high vascular density compared to less dense areas. Therefore, the proposed model can be extended to incorporate the effects of spatially heterogeneous blood vessel networks on intratumoral heterogeneity. In addition, we did not account for blood vessel generation and migration in response to angiogenic factors (e.g., VEGF, FGF) secreted from the hypoxic core of the tumor. Consequently, the current framework can be expanded into a spatiotemporal model that explicitly incorporates angiogenic signaling as a driver of vascularized tumor growth.

## Acknowledgment

SG gratefully acknowledges the Ministry of Education, Government of India, for research fellowship funding and the Indian Institute of Technology Guwahati for computational support.

## Conflict of interest

The authors declare no conflict of interest.

## A Appendix

### A.1 Hyper-vascularized tumor

Here we determine the analytical solution of our model when the tumor is fully vascularized, that is, *v → v_∞_*. From Eq. (2) the term 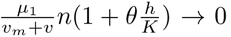, so, 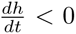. The rate of change of hypoxic cells is negative; that is, hypoxic cells slowly convert to normoxic cells, and eventually all hypoxic cells are converted to normoxic cells. At that point, *h* becomes 0. When the tumor is fully vascularized, the switching rate from normoxic to necrotic cells and the degradation of necrotic cells slow down. Therefore, *δ*_1_ = 0, *ν* = 0. Now, from Eq. (3), 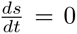, with the initial condition *s*(*t* = 0) = 0, necrotic cells becomes 0. Hence, Eq. (1) becomes

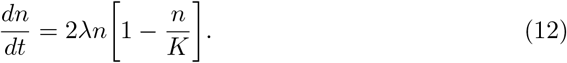

The solution of the Eq. (12) is given by

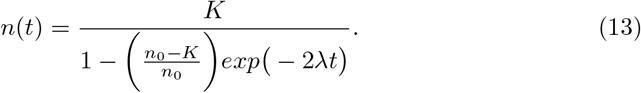

We take *v_∞_* = 130 to compute the numerical simulations. We observe that the numerical result is well agreement with the analytical counterpart (Figure 9).

### A.2 Hypo-vascularized tumor

Here, we find the analytical solution for a poorly vascularized or avascular tumor. For a poorly vascularized or avascular tumor *v →* 0. Now if *v_m_* is very small then the term 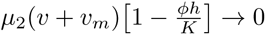. Recall Eq. (1)

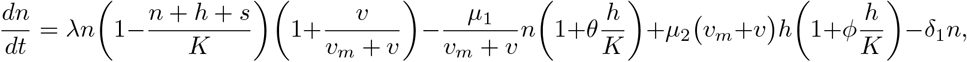

with the above assumptions, the equation becomes

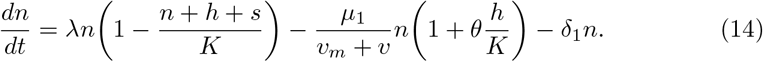

Since *v →* 0 and if *v_m_* is very small then, 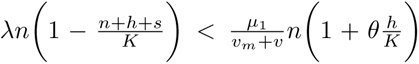.

Therefore 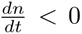. Eventually all normoxic cells are converted to hypoxic cells and after some time *n* becomes 0. Hence, Eq. (2) becomes

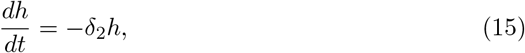

which gives the solution *h*(*t*) = *h*_1_*exp*(*−δ*_2_*t*), where initial conditions are taken as *n*(*t* = 0) = *n*_1_*, h*(*t* = 0) = *h*_1_*, s*(*t* = 0) = *s*_1_*, v*(*t* = 0) = *v*_1_. Now we recall Eq. (3)

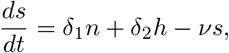

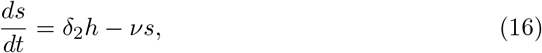

the solution of Eq. (16) is given by

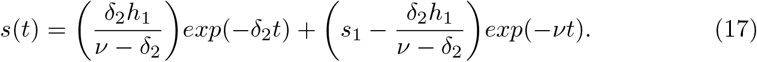

We can see that both *h*(*t*) and *s*(*t*) *→* 0 as *t → ∞*. So without vascularization tumor eventually fade away. Now if *ν* = 0 then

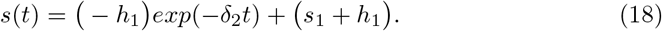

In that case only necrotic cells remains other cells fades as *t → ∞*.

## Notes

### Competing Interest Statement

The authors have declared no competing interest.

## References

[1] Hanahan, D., Weinberg, R.A.: The hallmarks of cancer. Cell 100(1), 57–70 (2000)

[2] Sadhu, G., Yadav, K., Ghosh, S.S., Dalal, D.: The impact of oxygen distribution on the tumor necrotic region: A two-phase model. Acta Biotheoretica 74(2), 9 (2026)

[3] Sadhu, G., Dalal, D.: A mathematical study of the interaction between oxygen and lactate in an in vivo and in vitro tumor. International Journal of Biomathematics, 2450138 (2024)

[4] Bull, J.A., Byrne, H.M.: The hallmarks of mathematical oncology. Proceedings of the IEEE 110(5), 523–540 (2022)

[5] Ribatti, D.: Judah folkman, a pioneer in the study of angiogenesis. Angiogenesis 11(1), 3–10 (2008)

[6] Breward, C.J.W., Byrne, H.M., Lewis, C.E.: A multiphase model describing vascular tumour growth. Bulletin of Mathematical Biology 65(4), 609–640 (2003)

[7] Owen, M.R., Alarćon, T., Maini, P.K., Byrne, H.M.: Angiogenesis and vascular remodelling in normal and cancerous tissues. Journal of mathematical biology 58(4), 689–721 (2009)

[8] Fiandaca, G., Bernardi, S., Scianna, M., Delitala, M.E.: A phenotype-structured model to reproduce the avascular growth of a tumor and its interaction with the surrounding environment. Journal of Theoretical Biology 535, 110980 (2022)

[9] Sadhu, G., Jain, P., George, J.T., Jolly, M.K.: A phenotype-structured pde framework for investigating the role of hypoxic memory on tumor invasion under cyclic hypoxia. Bulletin of Mathematical Biology 88(2), 23 (2026)

[10] Sadhu, G., Dalal, D.: Effects of non-linear interaction between oxygen and lactate on solid tumor growth under cyclic hypoxia. Bulletin of Mathematical Biology 87(3), 41 (2025)

[11] Yadav, K., Sadhu, G.: Effect of inosine on recurrence of tumor after radiation therapy: A mathematical investigation. Journal of Theoretical Biology, 112138 (2025)

[12] Sadhu, G.: A mathematical investigation of exosome and lactate levels interplay in an in vitro and in vivo tumors. Bulletin of Mathematical Biology 88(1), 4 (2026)

[13] Metzcar, J., Wang, Y., Heiland, R., Macklin, P.: A review of cell-based computational modeling in cancer biology. JCO clinical cancer informatics 2, 1–13 (2019)

[14] Ward, J.P., King, J.R.: Mathematical modelling of avascular-tumour growth. Mathematical Medicine and Biology: A Journal of the IMA 14(1), 39–69 (1997)

[15] Sherratt, J.A., Chaplain, M.A.: A new mathematical model for avascular tumour growth. Journal of mathematical biology 43(4), 291–312 (2001)

[16] Chaplain, M.A.: Mathematical modelling of angiogenesis. Journal of neurooncology 50(1), 37–51 (2000)

[17] Yang, H.M.: Mathematical modeling of solid cancer growth with angiogenesis. Theoretical Biology and Medical Modelling 9(1), 2 (2012)

[18] Breward, C., Byrne, H., Lewis, C.: Modelling the interactions between tumour cells and a blood vessel in a microenvironment within a vascular tumour. European Journal of Applied Mathematics 12(5), 529–556 (2001)

[19] Hubbard, M.E., Byrne, H.M.: Multiphase modelling of vascular tumour growth in two spatial dimensions. Journal of theoretical biology 316, 70–89 (2013)

[20] Breward, C.J., Byrne, H.M., Lewis, C.E.: The role of cell-cell interactions in a two-phase model for avascular tumour growth. Journal of mathematical biology 45(2), 125–152 (2002)

[21] Byrne, H.M., King, J.R., McElwain, D.S., Preziosi, L.: A two-phase model of solid tumour growth. Applied Mathematics Letters 16(4), 567–573 (2003)

[22] Swanson, K.R., Rockne, R.C., Claridge, J., Chaplain, M.A., Alvord Jr, E.C., Anderson, A.R.: Quantifying the role of angiogenesis in malignant progression of gliomas: in silico modeling integrates imaging and histology. Cancer research 71(24), 7366–7375 (2011)

[23] Stamper, I., Byrne, H., Owen, M., Maini, P.: Modelling the role of angiogenesis and vasculogenesis in solid tumour growth. Bulletin of mathematical biology 69(8), 2737–2772 (2007)

[24] Villa, C., Chaplain, M.A., Lorenzi, T.: Modeling the emergence of phenotypic heterogeneity in vascularized tumors. SIAM Journal on Applied Mathematics 81(2), 434–453 (2021)

[25] Borzouei, M., Mardaani, M., Emadi-Baygi, M., Rabani, H.: Development of a coupled modeling for tumor growth, angiogenesis, oxygen delivery, and phenotypic heterogeneity. Biomechanics and Modeling in Mechanobiology 22(3), 1067–1081 (2023)

[26] Sadhu, G., Byrne, H.M., Dalal, D.: The impact of oxygen heterogeneity on epithelial-mesenchymal transitions: a numerical study. Journal of Mathematical Biology 92(1), 15 (2026)

[27] Le, A., Stine, Z.E., Nguyen, C., Afzal, J., Sun, P., Hamaker, M., Siegel, N.M., Gouw, A.M., Kang, B.-h., Yu, S.-H., et al.: Tumorigenicity of hypoxic respiring cancer cells revealed by a hypoxia–cell cycle dual reporter. Proceedings of the National Academy of Sciences 111(34), 12486–12491 (2014)

[28] Forýs, U., Marciniak-Czochra, A.: Logistic equations in tumour growth modelling. International journal of applied mathematics and computer science 13(3), 317– 325 (2003)

[29] Durand, R.E., Raleigh, J.A.: Identification of nonproliferating but viable hypoxic tumor cells in vivo. Cancer research 58(16), 3547–3550 (1998)

[30] Druker, J., Wilson, J.W., Child, F., Shakir, D., Fasanya, T., Rocha, S.: Role of hypoxia in the control of the cell cycle. International Journal of Molecular Sciences 22(9), 4874 (2021)

[31] Goda, N., Ryan, H.E., Khadivi, B., McNulty, W., Rickert, R.C., Johnson, R.S.: Hypoxia-inducible factor 1*α* is essential for cell cycle arrest during hypoxia. Molecular and cellular biology 23(1), 359–369 (2003)

[32] Malekian, S., Rahmati, M., Sari, S., Kazemimanesh, M., Kheirbakhsh, R., Muhammadnejad, A., Amanpour, S.: Expression of diverse angiogenesis factor in different stages of the 4t1 tumor as a mouse model of triple-negative breast cancer. Advanced pharmaceutical bulletin 10(2), 323 (2020)

[33] Liu, W., Hillen, T., Freedman, H.: A mathematical model for m-phase specific chemotherapy including the g0-phase and immunoresponse. Mathematical Biosciences and Engineering 4(2), 239 (2007)

[34] Pinho, S.T.R.d., Bacelar, F.S., Andrade, R.F.S., Freedman, H.: A mathematical model for the effect of anti-angiogenic therapy in the treatment of cancer tumours by chemotherapy. Nonlinear Analysis: Real World Applications 14(1), 815–828 (2013)

[35] Sadhukhan, S., Basu, S.: Avascular tumour growth models based on anomalous diffusion: S. sadhukhan, sk basu. Journal of Biological Physics 46(1), 67–94 (2020)

[36] Zeng, Z., Zhao, Y., Chen, Q., Zhu, S., Niu, Y., Ye, Z., Hu, P., Chen, D., Xu, P., Chen, J., et al.: Hypoxic exosomal hif-1*α*-stabilizing circznf91 promotes chemoresistance of normoxic pancreatic cancer cells via enhancing glycolysis. Oncogene 40(36), 5505–5517 (2021)

[37] Godet, I., Oza, H.H., Shi, Y., Joe, N.S., Weinstein, A.G., Johnson, J., Considine, M., Talluri, S., Zhang, J., Xu, R., et al.: Hypoxia induces ros-resistant memory upon reoxygenation in vivo promoting metastasis in part via muc1-c. Nature communications 15(1), 8416 (2024)

[38] Baeriswyl, V., Christofori, G.: The angiogenic switch in carcinogenesis. In: Seminars in Cancer Biology, vol. 19, pp. 329–337 (2009). Elsevier

[39] Saltelli, A., Annoni, P., Azzini, I., Campolongo, F., Ratto, M., Tarantola, S.: Variance based sensitivity analysis of model output. design and estimator for the total sensitivity index. Computer physics communications 181(2), 259–270 (2010)

[40] Saltelli, A., Ratto, M., Andres, T., Campolongo, F., Cariboni, J., Gatelli, D., Saisana, M., Tarantola, S.: Global Sensitivity Analysis: the Primer. John Wiley & Sons, ??? (2008)

[41] Burrell, R.A., Swanton, C.: Tumour heterogeneity and the evolution of polyclonal drug resistance. Molecular oncology 8(6), 1095–1111 (2014)

[42] Ghajar, C.M., Peinado, H., Mori, H., Matei, I.R., Evason, K.J., Brazier, H., Almeida, D., Koller, A., Hajjar, K.A., Stainier, D.Y., et al.: The perivascular niche regulates breast tumour dormancy. Nature cell biology 15(7), 807–817 (2013)

